# Two species of *Aedes* show altitudinal variation in temperature preference in the islands of the Gulf of Guinea

**DOI:** 10.64898/2026.02.04.703755

**Authors:** Daniel R. Matute

## Abstract

**Background:** Temperature choice is a vector trait that influences microhabitat selection and can have important implications for vector species, as it may affect how often vectors encounter hosts. *Aedes aegypti* and *Ae. albopictus* are disease vectors whose geographic ranges continue to expand each year. One aspect that remains largely understudied is the altitudinal range of these species and the extent of differences in thermal behavior between lowland and highland populations.

**Methods:** I collected *Ae. aegypti* and *Ae. albopictus* on the islands of Bioko and São Tomé. I compared the distribution of the two species along an altitudinal cline spanning 2,000 m of elevation. I then used live specimens to test temperature preference for both species in a laboratory thermocline.

**Results:** I report the distribution of these two species on the island of Bioko and show that the abundance of immature stages of both species follows a negative exponential decay with altitude. I compare this distribution with that observed on the neighboring island of São Tomé, also in the Gulf of Guinea. Overall, the distribution patterns of the two species are similar, but models indicate a higher abundance at sea level in São Tomé than in Bioko. I used specimens from this survey to study temperature preference under controlled conditions. I found no significant differences between species or between sexes; however, I detected an altitudinal cline in temperature preference, with high-elevation populations preferring cooler temperatures on both islands.

**Conclusions:** These results indicate the presence of phenotypic variation in a key trait—temperature choice—that may alter the likelihood of contact between these vectors and humans.

## INTRODUCTION

Temperature plays a crucial role in shaping the physiological processes of insects, influencing their metabolism, development, and behavior (Gilbert and Raworth 1996; Bale 2002; Colinet et al. 2015; Bjørge et al. 2018; Mutamiswa et al. 2023). As ectothermic organisms, insects are particularly sensitive to environmental temperature, which strongly influences their internal physiological conditions (Tao et al. 2023). Variations in temperature can significantly affect enzymatic activities, metabolic rates, and life cycle events such as reproduction and growth (Matute et al. 2009; Abram et al. 2017; Brandt et al. 2018). Furthermore, temperature extremes can impose stress, leading to altered behaviors, reduced fitness, and increased mortality, ultimately determining the limits of geographic and ecological ranges (Bale 2002; González-Tokman et al. 2020). Understanding the relationship between temperature and insect physiology is essential for predicting insect population dynamics, with particular importance for disease vectors (Githeko et al. 2000; Gage et al. 2008; Ying Zhang et al. 2008; Sternberg and Thomas 2014; Caminade et al. 2019).

Among disease vectors, species of the genus *Aedes* have increased in importance over the last few decades. *Aedes aegypti* and *Aedes albopictus* are vectors for numerous arboviruses, including dengue, Zika, and chikungunya, among many other diseases (Gratz 2004; Jansen and Beebe 2010). Temperature modulates *Aedes* life cycle processes, including oviposition, larval development, and adult survival (Liu et al. 2017; Reinhold et al. 2018a). High temperatures typically accelerate development rates, reducing the duration of the aquatic phase (Bar-Zeev 1958; Couret et al. 2014; Couret and Benedict 2014). Extreme temperatures can reduce longevity (Brady et al. 2013), decrease survival, and induce morphological changes (Nik Abdull Halim et al. 2022), ultimately affecting population dynamics (Simoy et al. 2015). Since temperature impacts both the rate of maturation and the synchronization of life stages, temperature shifts can also influence vector emergence and the timing of disease outbreaks (Trejo et al. 2023). Understanding the ecological and physiological responses of *Aedes* mosquitoes to environmental temperatures is therefore critical for predicting their population dynamics, geographical distribution, and capacity to transmit pathogens, and for informing effective vector control strategies and public health interventions.

*Aedes aegypti* and *Aedes albopictus* occur on the islands of the Gulf of Guinea. *Aedes aegypti* is thought to be endemic to the African continent, including the islands of the Gulf of Guinea (Kamgang et al. 2024)*. Aedes albopictus* is a more recent introduction to the African continent and its islands. The first report of this species on the island of Bioko dates to 2001 (Toto et al. 2003) while the first report on the island of São Tomé dates to 2013 (Rader et al. 2024). On São Tomé, both of these *Aedes* species follow an altitudinal cline in abundance, being more abundant in the lowlands. However, the two species differ in their upper altitudinal boundaries. For both species, the upper altitudinal boundary of their range on São Tomé is between 1,500 and 2,000 m of elevation (Rader et al. 2024).

Among disease vectors, behavioral variation in traits that are key for habitat choice remains largely understudied. Studies of other disease-vector behavioral traits have nonetheless documented substantial variation. For example, *Anopheles arabiensis* individuals collected in human settlements in Tanzania were less likely to seek humans than cattle, whereas individuals from the same species collected in rural environments were much more likely to be attracted to humans (Mlacha et al. 2020). Habitat choice is one facet of insect biology that determines how often vectors come into contact with human hosts (Thongsripong et al. 2021). In *Aedes*, females are attracted to sites with higher human density (Rodrigues et al. 2015), which may represent an increased risk of disease transmission. Among the environmental factors that influence habitat choice, temperature is likely to play an important role. From a public health perspective, habitat choice across microclimatic conditions that increase contact with humans may directly influence disease risk (Githeko et al. 2000; Gage et al. 2008; Vonesch et al. 2016; Caminade et al. 2019). From an evolutionary perspective, the ability to seek permissive environments may be advantageous, promoting the evolution of behavioral syndromes that buffer populations from selection imposed by different thermal regimes and ultimately constrain physiological evolution. Quantifying phenotypic variability in habitat preference in disease vectors is therefore an important avenue for systematic study, and the altitudinal distribution of *Aedes* on the islands of the Gulf of Guinea lends itself well to such a survey.

In this study, I show that a behavioral trait—temperature choice—varies across populations on the African islands of São Tomé and Bioko. First, I document an altitudinal cline in the abundance of both *Ae. aegypti* and *Ae. albopictus* on the island of Bioko. High-elevation populations of both species show a stronger preference for lower temperatures than low-elevation populations, consistent with the cooler environmental conditions at higher elevations. I observe a similar pattern of temperature preference variation on the island of São Tomé.

Overall, both *Ae. aegypti* and *Ae. albopictus* show parallel patterns across the two altitudinal gradients. These results indicate that the sources of variation in behavioral traits are multifaceted and that, in disease vectors, understanding the drivers of phenotypic variation is crucial for assessing their potential to expand their geographic range.

## MATERIALS AND METHODS

### Sampling locations

I focused on two *Aedes* species with wide geographical distributions and high importance for disease transmission, *Ae. albopictus* and *Ae. aegypti* (Gratz 2004; Powell and Tabachnick 2013; Matthews 2019). To study abundance along the altitudinal gradient on Bioko, I sampled 14 locations along a transect starting in the city of Malabo (sea level) and extending to Lago Biao (1,944.6 m). I also sampled the summit of Pico Basile, the highest point on the island (2,825.1 m). In addition, I collected live specimens from four locations spanning over 1,000 m of elevation (27.2 m, 286.1 m, 339.7 m, and 1,064.4 m).

For São Tomé, collections used to assess altitudinal abundance have been reported elsewhere (Rader et al. 2024). For abundance studies in São Tomé, I only used the counts from 2013 to estimate the abundance distribution. I limited the analyses to this year because it was the same year for which I had collected on Bioko. I sampled four locations spanning approximately 1,000 m of elevation on the northern side of the island, between the settlements of São Tomé city and Montecafe (6.9 m, 292.3 m, 475.7 m, and 1,151.1 m). The number of specimens collected at each location is shown in Table S1.

### Specimen collection

Sampling procedures were similar across locations and followed two regimes targeting different life stages. Immature stages were collected from pools of standing water using stainless steel–coated mosquito dippers (John W. Hock Company, Gainesville, FL). Specimens were transferred to individual glass vials containing water. Vials were kept in a BugDorm (MegaView Science Co., Ltd., Taiwan), separated by species, to retain individuals until emergence.

To differentiate between *Ae. aegypti* and *Ae. albopictus* immature stages, I used non-destructive morphological traits. *Aedes aegypti* larvae possess comb scales on the terminal segment arranged in a single row of pitchfork-shaped scales, and the thorax shows prominent black hooks. In contrast, *Ae. albopictus* larvae have comb scales arranged in a single row of thorn-shaped scales and either small or absent thoracic hooks (Lee 1998). To differentiate pupae, I examined the terminal setae on the paddle: in *Ae. aegypti*, these setae are short and simple, whereas in *Ae. albopictus* they are longer and hair-like (Penn, 1949). Larvae and pupae were maintained in glass vials until adult emergence for further confirmation of species identity.

Adult mosquitoes were collected using a light trap composed of a white sheet and four light sources powered by 6 V batteries, deployed at sunset. Mosquitoes were aspirated as they landed and inspected morphologically (see below). Sampling was conducted for at least 10 days to obtain a minimum of 25 specimens per site. Adults were kept alive for subsequent experiments (see below).Table S1 summarizes the number of individuals collected.

### Temperature Preference Assay

I assessed temperature preference in both species using a thermocline constructed from a plexiglass chamber (dimensions: 12 cm × 45 cm × 1 cm) with an aluminum base. The design of the apparatus has been described previously (Matute et al. 2009; Cooper et al. 2018; Vivero-Gomez et al. 2024). A thermal gradient ranging from 18 °C to 30 °C was established using heat plates (120 VAC Thermo Scientific Cimarec Hot Plate, model UX-04600-01, Thermo Fisher Scientific, Waltham, MA, USA), increasing by approximately 2 °C every 6 cm across long axis of the chamber. While these temperatures are not considered extreme for either species (Reinhold et al. 2018a; Lahondère and Bonizzoni 2022), they reflect the range of temperatures in the islands of the Gulf of Guinea (Cooper et al. 2018; Comeault and Matute 2021). Each replicate included an average of 18 individuals (minimum = 4) captured adults. I did not include individuals collected as larvae to minimize the chance of assaying individuals from the same family. Individuals were allowed to move freely within the chamber for one hour. At the end of each trial, mosquitoes were isolated into seven partitions (10.5 cm × 6 cm × 1 cm) by sliding a rod connected to plexiglass dividers across the width of the chamber. I recorded the temperature in each partition using a Digi-Sense thermometer equipped with a type-T thermocouple (Cole-Parmer Instrument Co., Chicago, IL; catalog number 86460-05), as well as the number of individuals present in each partition. Males and females were evaluated separately to eliminate potential confounding effects of sexual attraction. Temperature preference of each sex was assessed in three independent replicates (i.e., the preference of an individual is only reported once). The number of individuals per replicate is listed in Table S2.

### Statistical analyses

I used the distributional and phenotypic data described above to study patterns of variation in abundance and thermal preference across the two transects for each *Aedes* species. All analyses were based on linear modeling approaches, which are described in detail below.

#### Abundance

I used abundance data to determine which functional form best explained the occurrence of *Aedes* species on Bioko. I fitted four families of models: one for immature stages and one for adult stages for each species. Each family comprised three functions: a linear model, an asymptotic model, and a negative exponential decay model. In all models, specimen abundance was the response variable, and altitude was the sole predictor. Linear models were fitted using the function *lm* (library *stats*, (Team 2023)) while exponential and asymptotic models were fitted using *nls.lm* (library *minpack.lm*, (Elzhov et al. 2016)). Table 1 describes the functional forms and parameters of each model. Model fit was compared using Akaike’s Information Criterion (AIC, (Akaike 1973)), and relative support was assessed using wAIC. In the majority of cases, the negative exponential decay function (See Results) was the best fit. To compare the parameters between the different functions, I generated bootstrap distributions (999 replicates) using the function *nlsBoot* (library (Baty et al. 2013)), which I then compared with the function *wilcox.test* (library stats, (Team 2023).

**TABLE 1.** Three models to differentiate to model the abundance of *Aedes* along an altitudinal cline on the islands of the Gulf of Guinea.

| Model | Function | Parameter explanation |
| --- | --- | --- |
| Linear | Abundance~ $a + b \times \text{altitude}$ | $a$: intercept<br>$b$: slope |
| Exponential | Abundance ~ $a \times e^{(-b \times \text{altitude})}$ | $a$: intercept<br>$b$: rate of decay |
| Asymptotic | Abundance ~ $d + (a - d) / (1 + (\text{altitude} / c)^b)$ | $a$: intercept<br>$b$: moment of change<br>$c$: rate of change<br>$d$: asymptote |

#### Temperature preference

To test whether altitude influenced temperature preference among populations from the Gulf of Guinea, I fitted linear models using the function *lm* (library *stats*, (Team 2023)), in which temperature choice (i.e., the partition in the thermocline) was the response variable, island, species, and sexwere included as fixed effects, and altitude was treated as a continuous predictor. The model also included the interactions between all effects and altitude. Differences in mean temperature preference between islands were assessed using Tukey’s Honest Significant Difference test, as implemented in the R function *ghlt* (library *multicomp*, (Hothorn et al. 2012; Hothorn et al. 2016)).

## RESULTS

### *Aedes* is more abundant in the lowlands than in the highlands of Bioko

In 2009, I found that *Ae. aegypti* and *Ae. albopictus* were present in the Bioko cities of Malabo, Luba, and Riaba; however, these collections did not allow me to determine the altitudinal distribution of the species because all three cities are located at sea level. These observations were consistent with previous studies reporting that the island harbors both *Ae. aegypti* and *Ae. albopictus* ((Giger et al. 2024) and (Toto et al. 2003), respectively). Four years later, in 2013, I expanded my sampling to six sites along an altitudinal gradient on Bioko to determine the upper elevational limits of the two species. Figure 1 shows the distributions of immature stages of *Ae. aegypti* (Figure 1A) and *Ae. albopictus* (Figure 1B) along this gradient. Figure 2 shows the distributions of adults along the same gradient. For both species, the upper elevational limit for adults was close to 1,500 m, with the exception of a single *Ae. albopictus* collection at nearly 2,000 m.

**FIGURE 1.**
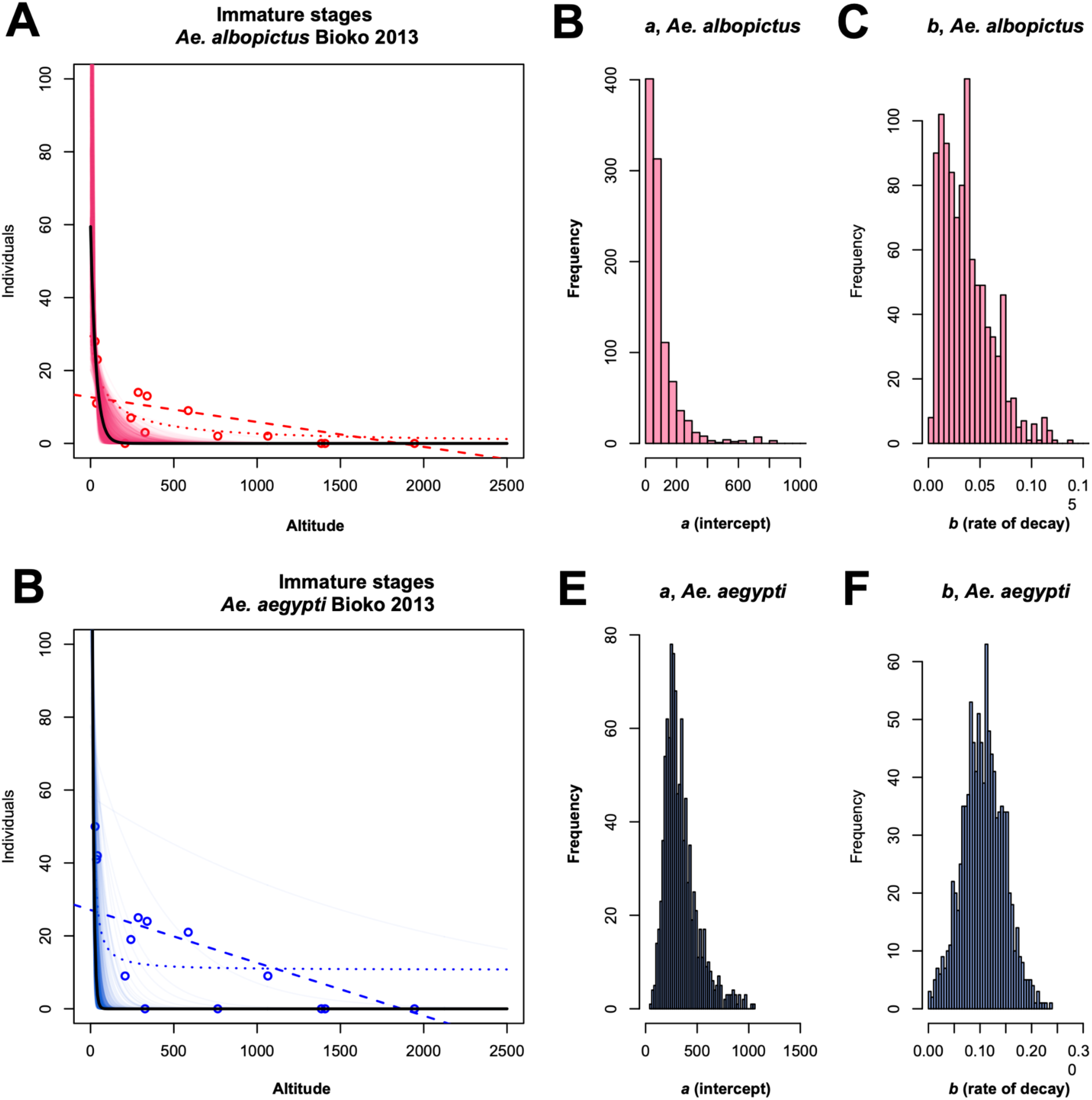
Abundance of immature individuals of two species of *Aedes* along an altitudinal gradient on the island of Bioko. **A.** Best-fitting model for immature stages of *Ae. albopictus*. The semitransparent lines represent 1,000 bootstrapped distributions for the best-fitting model, an exponential decay. **B.** Bootstrapped distributions of the intercept for the abundance function of immature stages of *Ae. albopictus*. **C.** Bootstrapped distributions of the inflexion point for the abundance function of immature stages of *Ae. albopictus*. **D.** Best fitting model for immature stages of *Ae. aegypti*. The semitransparent lines represent 1,000 bootstrapped distributions for the best-fitting model, an exponential decay. **E.** Bootstrapped distributions of the intercept for the abundance function of immature stages of *Ae. aegypti*. **F.** Bootstrapped distributions of the inflexion point for the abundance function of immature stages of *Ae. aegypti*.

**FIGURE 2.**
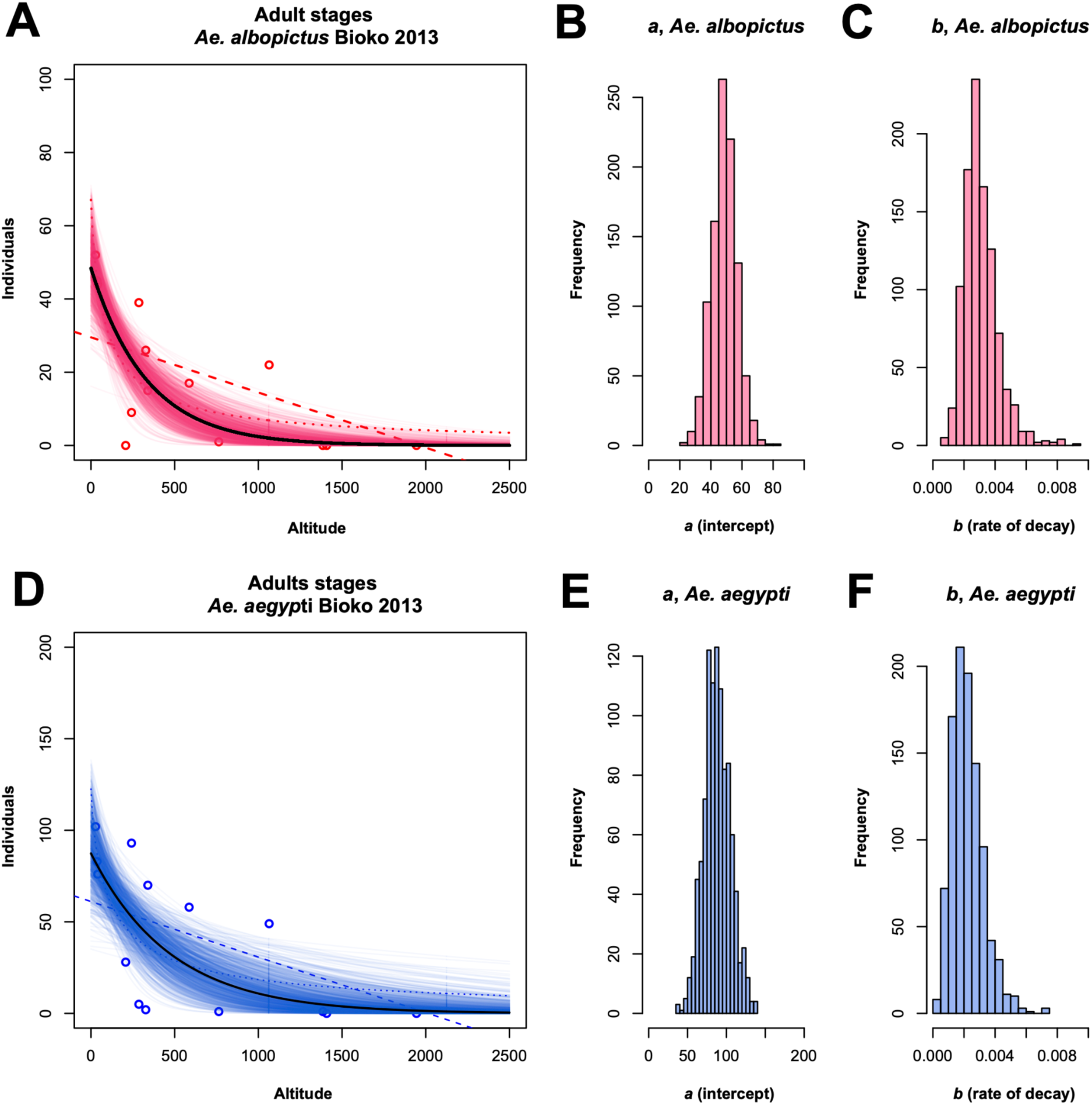
Abundance of adult individuals of two species of *Aedes* along an altitudinal gradient on the island of Bioko. **A.** Best-fitting model for adults of *Ae. albopictus*. The semitransparent lines represent 1,000 bootstrapped distributions for the best-fitting model, an exponential decay. **B.** Best-fitting model for adults of *Ae. aegypti*. The semitransparent lines represent 1,000 bootstrapped distributions for the best-fitting model, an exponential decay. **C.** Bootstrapped distributions of the intercept for the abundance function of adults of *Ae. albopictus*. **D.** Bootstrapped distributions of the inflection point for the abundance function of adults of *Ae. albopictus*. **E.** Bootstrapped distributions of the intercept for the abundance function of adults of *Ae. aegypti*. **F.** Bootstrapped distributions of the inflection point for the abundance function of adults of *Ae. aegypti*.

**FIGURE 3.**
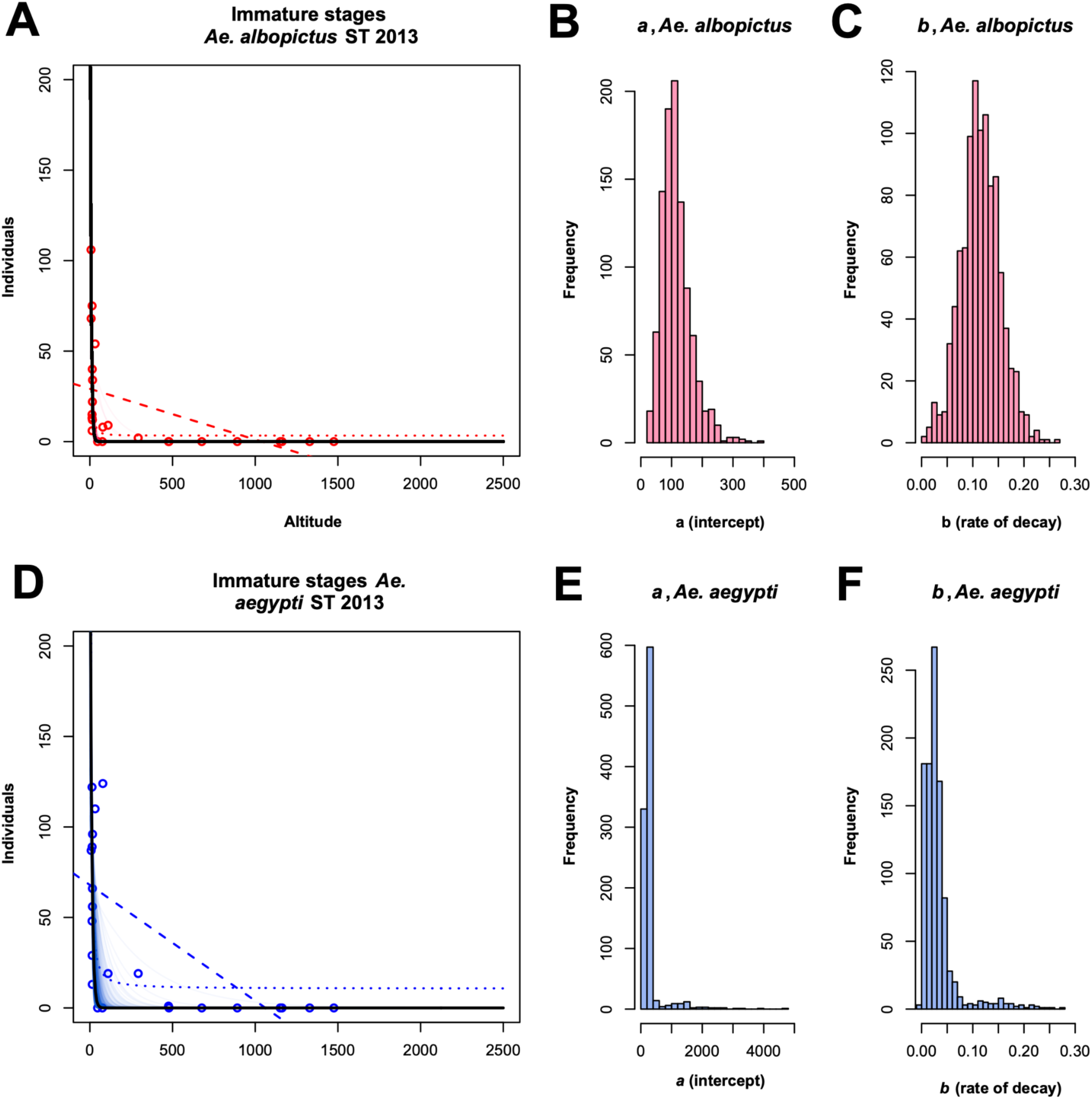
Abundance of immature individuals of two species of *Aedes* along an altitudinal gradient on the island of São Tomé. **A.** Best-fitting model for immature stages of *Ae. albopictus*. The semitransparent lines represent 1,000 bootstrapped distributions for the best-fitting model, an exponential decay. **B.** Best-fitting model for immature stages of *Ae. aegypti*. The semitransparent lines represent 1,000 bootstrapped distributions for the best-fitting model, an exponential decay. **C.** Bootstrapped distributions of the intercept for the abundance function of immature stages of *Ae. albopictus*. **D.** Bootstrapped distributions of the inflexion point for the abundance function of immature stages of *Ae. albopictus*. **E.** Bootstrapped distributions of the intercept for the abundance function of immature stages of *Ae. aegypti*. **F.** Bootstrapped distributions of the inflexion point for the abundance function of immature stages of *Ae. aegypti*.

**FIGURE 4.**
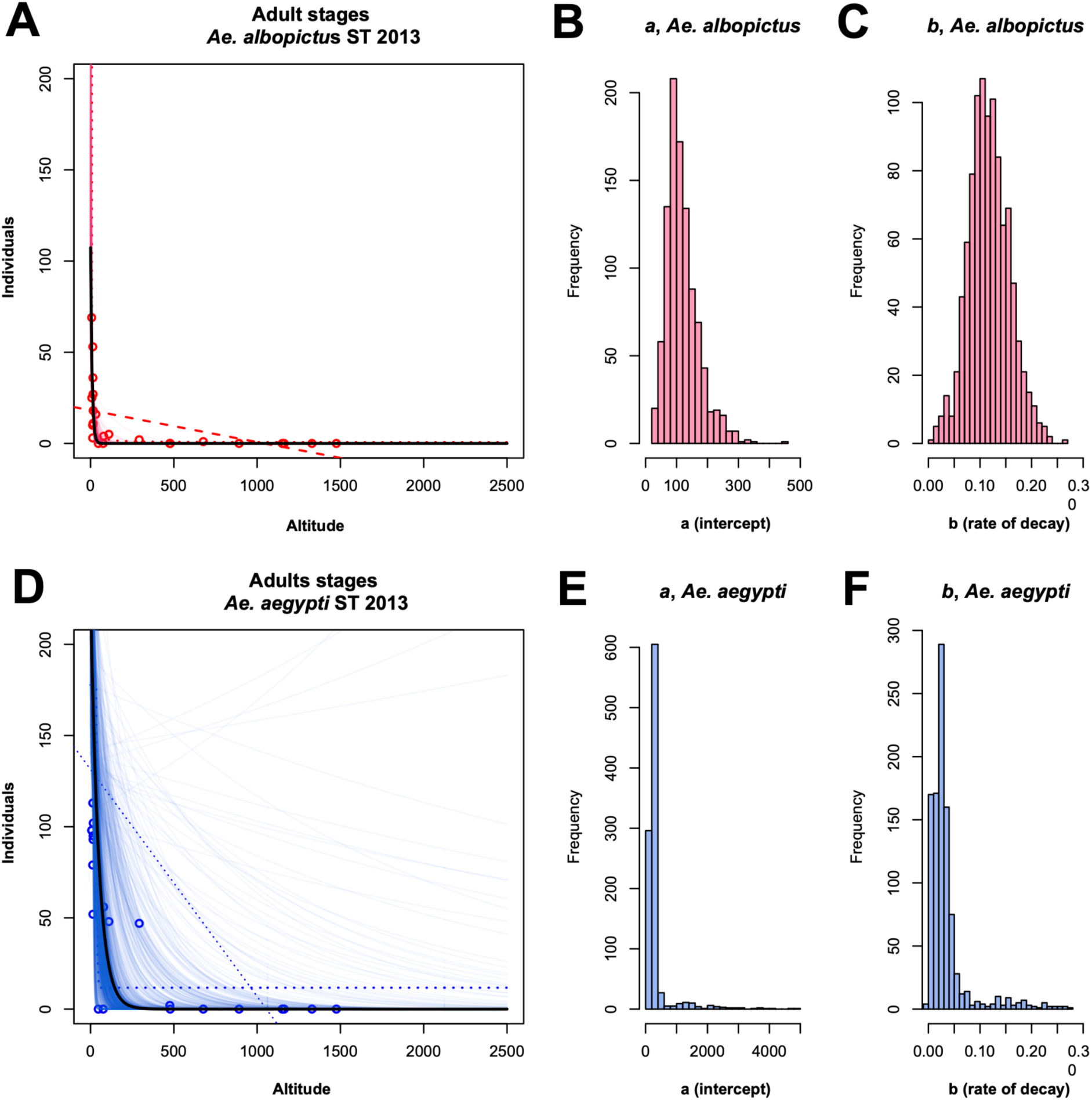
Abundance of adult individuals of two species of *Aedes* along an altitudinal gradient on the island of São Tomé. **A.** Best-fitting model for adults of *Ae. albopictus*. The semitransparent lines represent 1,000 bootstrapped distributions for the best-fitting model, an exponential decay. **B.** Best fitting model for adults of *Ae. aegypti*. The semitransparent lines represent 1,000 bootstrapped distributions for the best-fitting model, an exponential decay. **C.** Bootstrapped distributions of the intercept for the abundance function of adults of *Ae. albopictus*. **D.** Bootstrapped distributions of the inflection point for the abundance function of adults of *Ae. albopictus*. **E.** Bootstrapped distributions of the intercept for the abundance function of adults of *Ae. aegypti*. **F.** Bootstrapped distributions of the inflection point for the abundance function of adults of *Ae. aegypti*.

Using this distributional data, I fitted models to determine the function that best explained the distributions of the two species for both collection regimes (immature stages and adults). In three of the four cases, the negative exponential decay model provided a better fit than either the asymptotic or linear models (Table 2). The only exception was the distribution of *Ae. albopictus* immature stages, which was better explained by an asymptotic model. Because the biological interpretations of the asymptotic and exponential models are not categorically different—both assume a faster-than-linear decline in abundance with altitude—I used the exponential models to compare the rates of decay in abundance between the two *Aedes* species.

**TABLE 2.**
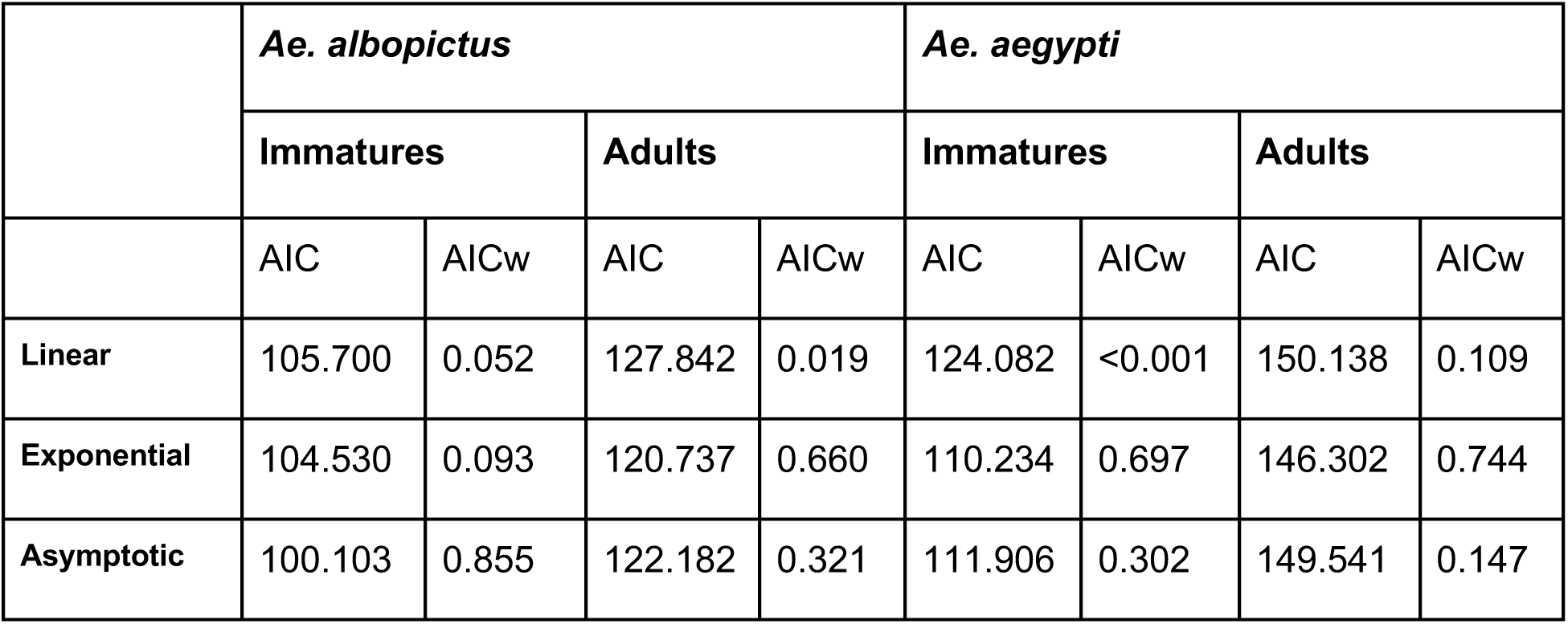
Model fitting for the distribution of *Ae. albopictus* and *Ae. aegypti* on the island of Bioko. AIC: Akaike Information Criterion. AICw: AIC weight.

|  | *Ae. albopictus* |  |  |  | *Ae. aegypti* |  |  |  |
| --- | --- | --- | --- | --- | --- | --- | --- | --- |
|  | Immatures |  | Adults |  | Immatures |  | Adults |  |
|  | AIC | AICw | AIC | AICw | AIC | AICw | AIC | AICw |
| Linear | 105.700 | 0.052 | 127.842 | 0.019 | 124.082 | <0.001 | 150.138 | 0.109 |
| Exponential | 104.530 | 0.093 | 120.737 | 0.660 | 110.234 | 0.697 | 146.302 | 0.744 |
| Asymptotic | 100.103 | 0.855 | 122.182 | 0.321 | 111.906 | 0.302 | 149.541 | 0.147 |

**TABLE 3.**
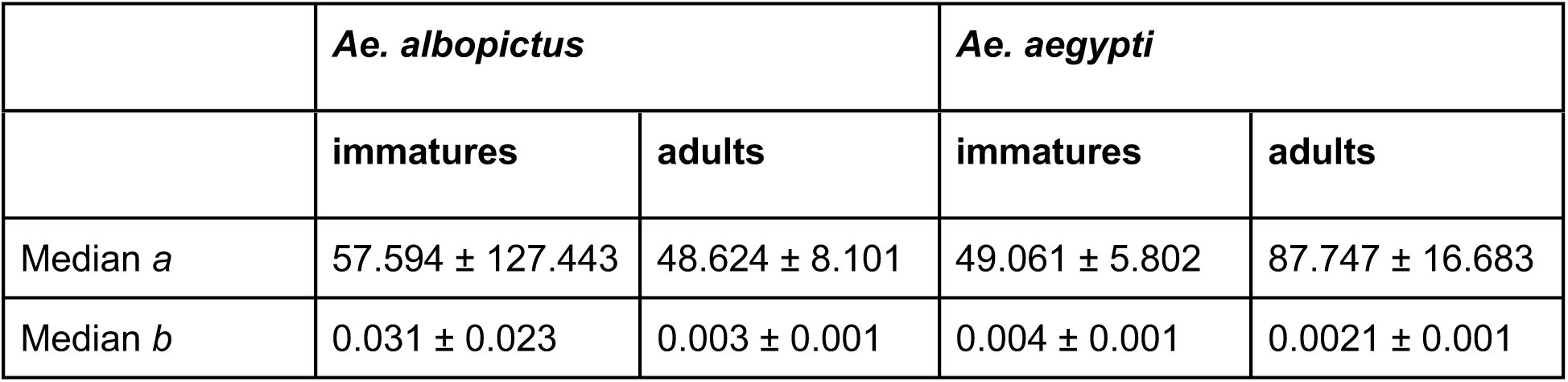
Intercept and rate of decay values for the negative exponential functions relating *Aedes* abundance and altitude on the island of Bioko. Note that the values are median estimates from the bootstrap functions shown in Figure 2.

First, I tested whether the rate of decay differed between developmental stages within species by comparing parameters of the exponential models. The intercept (representing inferred abundance at sea level) was higher for immature stages than for adults in *Ae. albopictus* (*P* < 0.001), whereas in *Ae. aegypti,* the intercept was higher for adults than for immatures (*P* < 0.001). For both species, the rate of decay was steeper for immature stages than for adults (*P* < 0.001). Next, I compared the parameters of these distributions between species for each developmental stage. For immature stages, both the intercept (*P* < 0.001) and the rate of decay (*P* < 0.001) differed between species. The same pattern was observed for adults, for which both the intercept (*P* < 0.001) and the rate of decay (*P* < 0.001) also differed between species. Table S3 lists all the pairwise comparisons.

### The abundance distributions of *Aedes* on Bioko and São Tomé are qualitatively similar, but quantitatively distinct

*Aedes albopictus* and *Ae. aegypti* are also present on the island of São Tomé (Reis et al. 2017; Kamgang et al. 2024). I used the previously published data on the distribution of the two *Aedes* on São Tomé (Rader et al. 2024) to determine the best-fitting function of their distribution in that island. As expected, and similar to Bioko, the highest abundance of the two species occurs at sea level. The upper boundary of *Ae. albopictus* was 676.8 m in 2013, but the species could be found at 1,150 m by 2023. *Aedes aegypti* followed a similar pattern: in 2013, the upper boundary was 475 m; by 2023, the species could be found at 1,162 m (Rader et al. 2024). In São Tomé, the distribution of the two species follows a similar pattern from Bioko as the exponential decay function is the best fit for both species’ abundance (Table 4). Similar to the results in Bioko, the intercepts differed between the functions for different developmental stages, and between species (W > 1385, *P <* 0.001, in all cases). These results indicate that, similar to Bioko, the distribution of *Ae. albopictus* and *Ae. aegypti* on the island of São Tomé shows differences in the parameters that best explain their distribution.

**TABLE 4.** Model fitting for the distribution of *Ae. albopictus* and *Ae. aegypti* on the island of São Tomé. AIC: Akaike Information Criterion. AICw: AIC weight.

|  | *Ae. albopictus* |  |  |  | *Ae. aegypti* |  |  |  |
| --- | --- | --- | --- | --- | --- | --- | --- | --- |
|  | Immatures |  | Adults |  | Immatures |  | Adults |  |
|  | AIC | AICw | AIC | AICw | AIC | AICw | AIC | AICw |
| Linear | 228.726 | <0.001 | 206.632 | <0.001 | 258.499 | 0.031 | 313.582 | 0.151 |
| Exponential | 210.541 | 0.577 | 190.927 | 0.739 | 253.455 | 0.389 | 310.508 | 0.701 |
| Asymptotic | 211.164 | 0.423 | 193.012 | 0.261 | 252.657 | 0.580 | 313.612 | 0.148 |

**TABLE 5.** Intercept and rate of decay values for the negative exponential functions relating *Aedes* abundance and altitude on the island of São Tomé. Note that the values are median estimates from the bootstrap functions shown in Figure 2.

|  | *Ae. albopictus* |  | *Ae. aegypti* |  |
| --- | --- | --- | --- | --- |
|  | Immatures | Adults | Immatures | Adults |
| Median *a* | 289.968 ± 127.762 | 108.384 ± 51.877 | 296.655 ± 169.284 | 233.538 ± 468.080 |
| Median *b* | 0.169±0.039 | 0.117 ± 0.039 | 0.104 ±0.041 | 0.025 ± 0.043 |

Finally, I compare the parameters of the abundance function between islands. The intercepts are higher in São Tomé than in Bioko for the four types of collections (*P <* 0.001 in all cases, Table S3). Similarly, the rate of decay is much higher for the transect in Bioko, which is not surprising given that both islands show no collected specimens at altitudes over 2,000m.

### Temperature preference follows an altitudinal cline

I studied the extent of phenotypic variation in temperature choice along the two altitudinal gradients on two islands within the Gulf of Guinea, Bioko and São Tomé. I found variation in temperature preference in both altitudinal gradients for both species. Figure 5 shows the mean temperature preferences along the two altitudinal gradients. Table 6 shows the result for a linear model to dissect the sources of variance in heterogeneity in temperature choice. The first notable result is that the temperature preference did not differ between *Ae. aegypti* and *Ae. albopictus*. The two species show a similar mean temperature preference and standard deviations (Preference*_Ae. aegypti_*= 21.953±3.705, Preference*_Ae. albopictus_*= 22.050±3.639). Neither the two- or three-way interactions involving species were significant either, thus suggesting that temperature preference is similar in these two species of *Aedes*, at least in the islands of the Gulf of Guinea.

**FIGURE 5.**
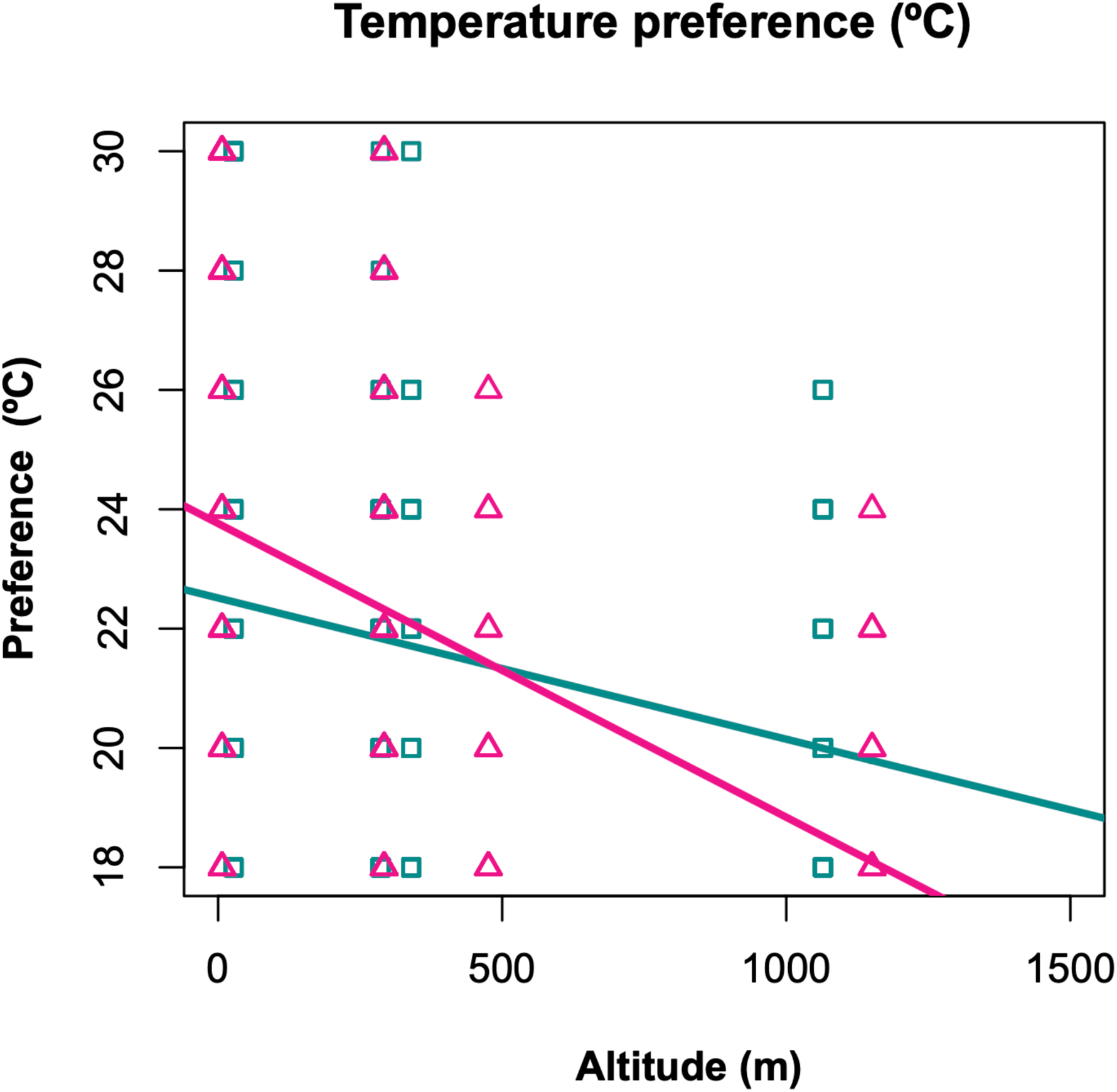
Temperature preference along the altitudinal gradient on the islands of Bioko and São Tomé. Pink: São Tomé, cyan: Bioko.

**TABLE 6.**
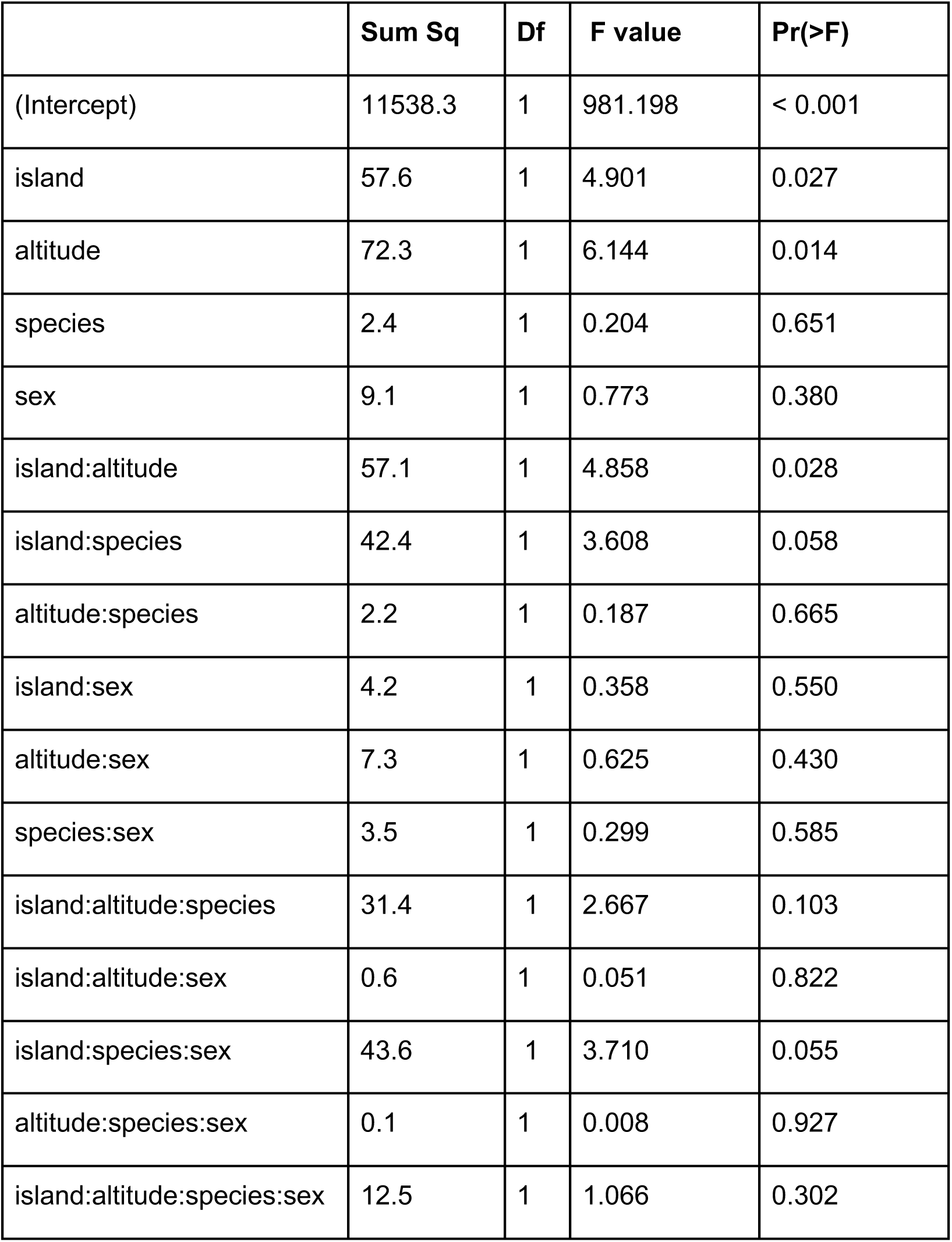
Temperature preference shows clinal variation on Bioko and São Tomé. A factorial linear model revealed heterogeneity in temperature preference across *Aedes* populations explained by island of origin, altitude and the interactions between these two factors. The df of the residuals was 7055.6.

The largest source of variation in the trait was altitude, which showed a negative relationship with temperature preference, indicating that populations from higher elevations prefer cooler temperatures. On average, temperature preference differed among populations that were separated by 100m of elevation by 0.271°C (SE=0.001, t= −2.479, *P* = 0.014).

Second, I detected significant differences in temperature preference depending on the island of origin. The temperature preference of populations from São Tomé is slightly higher than that of populations on the island of Bioko (Estimate=1.970, SE=0.890, t = 2.214, *P* = 0.027). These results indicate that the thermal behavior in *Aedes* is contingent on the island of origin. Finally, I detected a significant interaction between island and altitude, indicating that the relationships between altitude and temperature preference, albeit negative in both cases, differ between the two islands.

Sex was not a significant factor in the linear model, indicating no major differences in temperature choice between males and females; none of the interactions between sex and any other factor were significant. None of the three-way interactions, nor the four-way interaction, were significant.

## DISCUSSION

Thermal preference in mosquitoes plays a critical role in shaping their behavior, physiology, and ecological interactions (Reinhold et al. 2018b; Abbasi 2025). Some species of mosquitoes show distinct thermal preferences that influence habitat selection, feeding behavior, and reproductive success (Reinhold et al. 2022; Hug et al. 2023). These preferences can directly impact vectorial capacity, as temperature affects the replication rates of pathogens such as *Plasmodium* and arboviruses within the mosquito (Noden et al. 1995; Thu et al. 1998). In warmer and permissive environments, shorter extrinsic incubation periods may enhance transmission rates, whereas extreme temperature conditions may limit vector activity or pathogen development (Paaijmans et al. 2012; Winokur et al. 2020). In this report, I describe altitudinal variation in abundance for two *Aedes* species on the island of Bioko. Moreover, I compare these distributions with patterns of individual abundance across a similar altitudinal transect on the nearby island of São Tomé, also in the Gulf of Guinea. I also used a subset of specimens to measure the extent of thermal preference variation in populations of *Aedes aegypti* and *Ae. albopictus* on these two islands. I found differences among populations that can be explained by island of origin and altitude, but not by sex or species. In the following paragraphs, I discuss the implications of these findings.

The patterns of individual abundance observed on Bioko follow the same general trends reported in other altitudinal gradients, where vector species become rarer at higher elevations. For example, a survey in central Italy (Romiti et al. 2022) revealed that the best-fitting model for vector distribution was an exponential decay, similar to what I observed on Bioko and São Tomé. Although central Italy does not harbor *Ae. aegypti*, the upper altitude of occurrence for *Ae. albopictus* is approximately 600 m, with populations predicted to exist up to 1,015 m, posing challenges for surveillance at higher elevations than those currently monitored. While I report the presence of *Aedes* individuals at elevations close to 1,500 m, these records do not reach the elevations reported from Colombia (2,600m, (Gomez et al. 2025)) and Bolivia (2,500m, (Paupy et al. 2012; Ríos et al. 2023). Extensive modeling work has shown that the pace of geographic range expansion in *Aedes* is rapid (Gubler 2003; Kraemer et al. 2019; Longbottom et al. 2023), yet comparatively little attention has been paid to invasion along altitudinal gradients. In Italy, *Ae. albopictus* was first reported in 1992 (Dalla Pozza and Majori 1992), predating reports from the Gulf of Guinea by over a decade. Different populations of invasive species vary in their ability to adapt to new environments (e.g., (Kent et al. 2024)), but systematic assessments of this variation remain scarce. Longitudinal studies of expansion rates are therefore needed (e.g., (Romiti et al. 2022; Rader et al. 2024) to determine whether human populations at mid and high elevations will increasingly come into contact with these vectors and the diseases they transmit.

Beyond abundance, systematic collections combined with phenotypic characterization across a species’ geographic range can provide insight into the microhabitats different species choose and, consequently, their likelihood of contact with humans. Here, I show that temperature preference in two *Aedes* species decreases with altitude and differs between islands. These differences in temperature preference partly reflect variation in environmental temperatures along altitudinal gradients on São Tomé and Bioko ((Comeault and Matute 2021) and (Cooper et al. 2018) respectively), where higher elevations are consistently cooler than lower elevations. The results indicate geographic variation in this behavioral trait in both species, but no detectable differences between the two *Aedes* species. Similarly, a previous comparison between *Ae. aegypti* and *Ae. japonicus* found no differences in thermal preference along a laboratory thermal gradient in colonies maintained from New Caledonia (Verhulst et al. 2020). A fully phylogenetically informed survey will be required to determine the extent of interspecific differences in thermal behavior across this clade of disease vectors.

Several factors not explored here may also influence temperature choice. Infection with disease agents can alter thermal preference in unpredictable ways. *Aedes aegypti* experimentally infected with Sindbis virus preferred temperatures approximately 5 °C warmer than uninfected individuals, whereas *Ae. japonicus* experimentally infected with *Dirofilaria immitis* preferred temperatures about 4 °C cooler than non-infected individuals (Hug, Gretener-Ziegler, et al. 2024). Although the experimental setup using a thermal gradient on an aluminum plate allowed precise control of temperature, other environmental and internal factors—such as relative humidity, shade, and blood-feeding status—are also likely to influence *Aedes* resting place choice ((Ziegler et al. 2023) but see (Verhulst et al. 2020)).

This study also highlights several avenues for future research. While I characterized thermal behavior, I did not measure thermal limits, precluding an assessment of whether behavioral and physiological components of thermal fitness are integrated. Moreover, altitudinal variation represents only one of several spatiotemporal factors that may generate and maintain phenotypic variation. Seasonal and latitudinal gradients were not examined here. Studies that integrate altitude, latitude, and seasonality would allow assessment of the relative contributions of these factors to behavioral variation in disease vectors. Few such comprehensive surveys exist; for example, Romiti et al. (Romiti et al. 2022) examined the combined effects of altitude and seasonality on *Ae. albopictus* abundance in central Italy. Integrating the multiple facets of geographic range (altitude, latitude), and seasonality will reveal how phenotypic and genetic variation are compartmentalized in disease vectors.

Climate change is expected to alter the geographic range and seasonal activity of *Aedes* mosquitoes. Both *Ae. aegypti* and *Ae. albopictus* have expanded their ranges at an alarming rate (Kraemer et al. 2019; Longbottom et al. 2023), and experimental evolution studies have demonstrated that thermal behavior can evolve rapidly, on the order of tens of generations (Hug, Kropf, et al. 2024). Rising global temperatures may facilitate range expansion into temperate regions while simultaneously increasing exposure to thermal extremes. In addition, urban heat islands—localized areas with elevated temperatures—may create microhabitats that either enhance or constrain mosquito proliferation, depending on species-specific thermal tolerances and preferences. Understanding temperature preferences is therefore critical for predicting future distributional shifts and assessing the risk of vector-borne diseases in newly colonized areas.

Together, these results indicate that both *Ae. aegypti* and *Ae. albopictus* are currently constrained to low elevations on Bioko, with abundance declining nonlinearly with altitude. Although both species can occur at relatively high elevations, their distributions are best characterized by rapid exponential decay, particularly for immature stages. Differences between species and developmental stages in the abundance function parameters suggest that *Ae. albopictus* and *Ae. aegypti* respond differently to environmental gradients on each island. While the human population densities at higher elevations are sparse in both islands, these results have implications for how invasive species might move along altitudinal gradients of densely-populated centers (e.g., the Andes and the Himalayas). These patterns are consistent with the idea that abiotic factors associated with elevation, such as temperature, play a key role in shaping *Aedes* distributions and may limit the potential for sustained highland populations.

## Acknowledgements

I would like to thank the Matute lab for helpful comments. DRM was supported by the National Institute of General Medical Sciences under Award R35GM148244.

## SUPPLEMENTARY TABLES

**TABLE S1.** Number of specimens from two different species of *Aedes* collected on São Tomé and Bioko. ST: São Tomé.

|  |  |  | *Ae. albopictus* |  | *Ae. aegypti* |  |
| --- | --- | --- | --- | --- | --- | --- |
| Island | Location | Altitude | Immatures | Adults | Immatures | Adults |
| Bioko | Riaba | 27.2 | 28 | 52 | 50 | 102 |
| Bioko | Luba1 | 36 | 11 | 45 | 41 | 83 |
| Bioko | Ela Nguema | 40.5 | 23 | 44 | 42 | 76 |
| Bioko | Luba6 | 207.2 | 0 | 0 | 9 | 28 |
| Bioko | Bakake | 242.1 | 7 | 9 | 19 | 93 |
| Bioko | Rebola | 286.1 | 14 | 39 | 25 | 5 |
| Bioko | Rilaja | 327.1 | 3 | 26 | 0 | 2 |
| Bioko | Road6 | 339.7 | 13 | 15 | 24 | 70 |
| Bioko | Road5 | 586.4 | 9 | 17 | 21 | 58 |
| Bioko | Road4 | 763.7 | 2 | 1 | 0 | 1 |
| Bioko | Loreto | 1064.4 | 2 | 22 | 9 | 49 |
| Bioko | Cascade | 1385.5 | 0 | 0 | 0 | 1 |
| Bioko | Nova Moka | 1408 | 0 | 0 | 0 | 0 |
| Bioko | Lago Niao | 1944.6 | 0 | 0 | 0 | 0 |
| Bioko | Basile | 2825.1 | 0 | 0 | 0 | 0 |
| ST | Airport1 | 6.9 | 106 | 69 | 214 | 98 |
| ST | Pestana | 7.7 | 68 | 25 | 87 | 98 |
| ST | Lagarto | 12.9 | 15 | 3 | 122 | 783 |
| ST | Morro Peixe | 12.9 | 13 | 10 | 48 | 79 |
| ST | Neves | 13.3 | 75 | 10 | 89 | 228 |
| ST | Sweet_GH | 13.4 | 6 | 53 | 29 | 113 |
| ST | Praia_grande<br>_1 | 14.4 | 40 | 36 | 13 | 52 |
| ST | Santana2 | 15.1 | 12 | 11 | 56 | 93 |
| ST | Porto Alegre | 15.2 | 22 | 27 | 66 | 95 |
| ST | Grelados | 16 | 34 | 18 | 96 | 102 |
| ST | Santo Amaro<br>2 | 31.5 | 54 | 16 | 110 | 214 |
| ST | Santo Amaro | 46.7 | 0 | 0 | 0 | 0 |
| ST | Santana | 75.5 | 0 | 0 | 0 | 0 |
| ST | Caue | 78.3 | 8 | 4 | 124 | 56 |
| ST | Bobo Forro | 109.9 | 9 | 5 | 19 | 48 |
| ST | Trindade | 292.3 | 2 | 2 | 19 | 47 |
| ST | Batepa1 | 475.7 | 0 | 0 | 1 | 2 |
| ST | Batepa2 | 478.3 | 0 | 0 | 0 | 0 |
| ST | Monte Cafe | 676.8 | 0 | 1 | 0 | 0 |
| ST | San Nicolau | 891.6 | 0 | 0 | 0 | 0 |
| ST | Bom Suceso | 1151.1 | 0 | 0 | 0 | 0 |
| ST | Trap10 | 1162.1 | 0 | 0 | 0 | 0 |
| ST | Fork | 1329.2 | 0 | 0 | 0 | 0 |
| ST | Lago Amelia | 1475.6 | 0 | 0 | 0 | 0 |

**TABLE S2.** Number of individuals per replicate of the behavioral assay.

| Island | Location name | Altitude | Species | Number of individuals |
| --- | --- | --- | --- | --- |
| ST | airport1 | 6.9 | *Ae. albopictus* | 56 |
| ST | airport1 | 6.9 | *Ae. aegypti* | 56 |
| ST | Trindade | 292.3 | *Ae. albopictus* | 67 |
| ST | Trindade | 292.3 | *Ae. aegypti* | 74 |
| ST | Batepa | 475.7 | *Ae. albopictus* | 16 |
| ST | Batepa | 475.7 | *Ae. aegypti* | 12 |
| Bioko | Riaba | 27.2 | *Ae. albopictus* | 90 |
| **Bioko** | Riaba | 27.2 | *Ae. aegypti* | 0 |
| **Bioko** | Rebola | 286.1 | *Ae. albopictus* | 44 |
| **Bioko** | Rebola | 286.1 | *Ae. aegypti* | 21 |
| **Bioko** | road_6 | 339.7 | *Ae. albopictus* | 21 |
| **Bioko** | road_6 | 339.7 | *Ae. aegypti* | 0 |
| **Bioko** | Loreto | 1064.4 | *Ae. albopictus* | 23 |
| **Bioko** | Loreto | 1064.4 | *Ae. aegypti* | 47 |

### Minimium 4

**TABLE S3.**
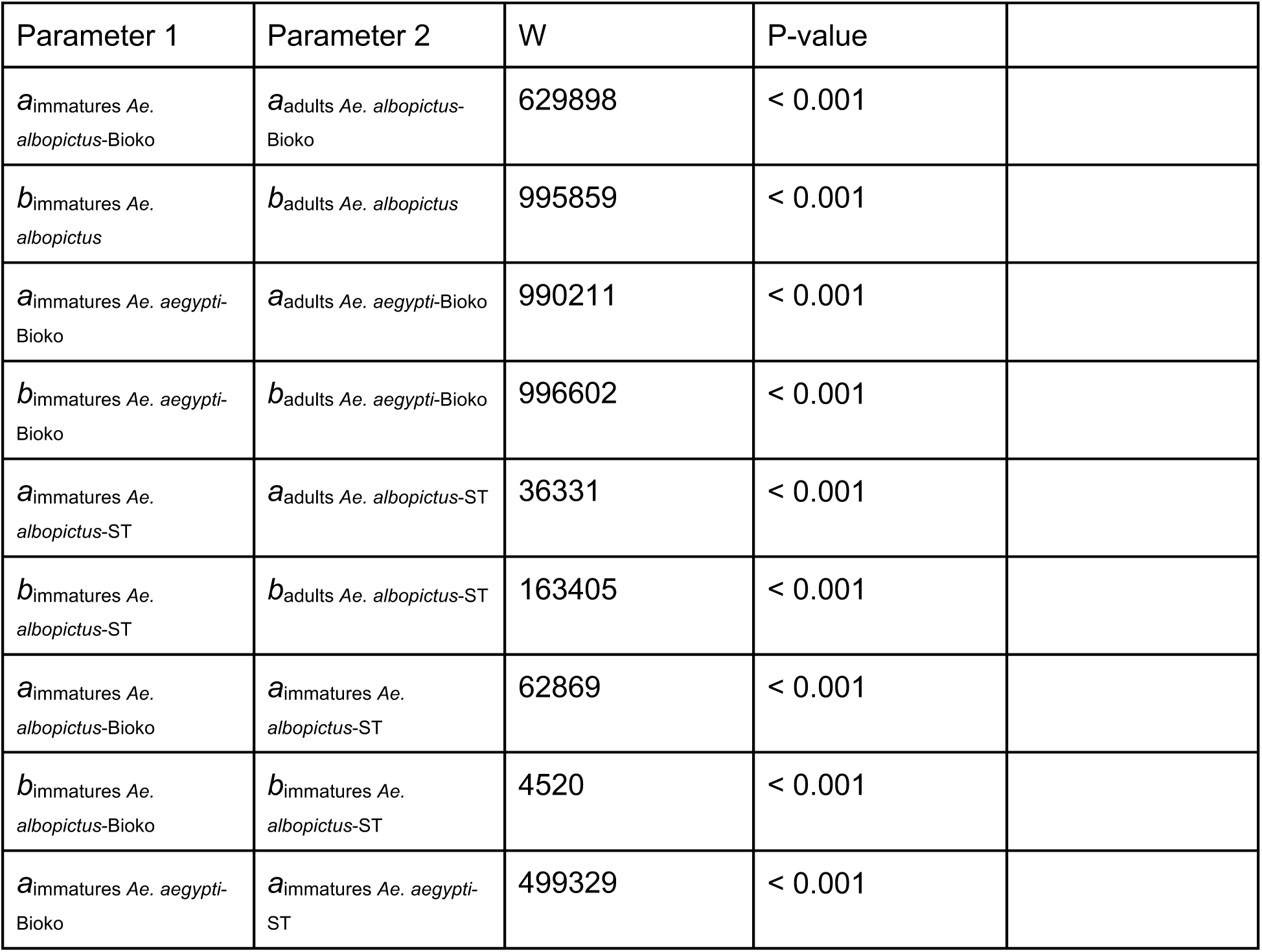
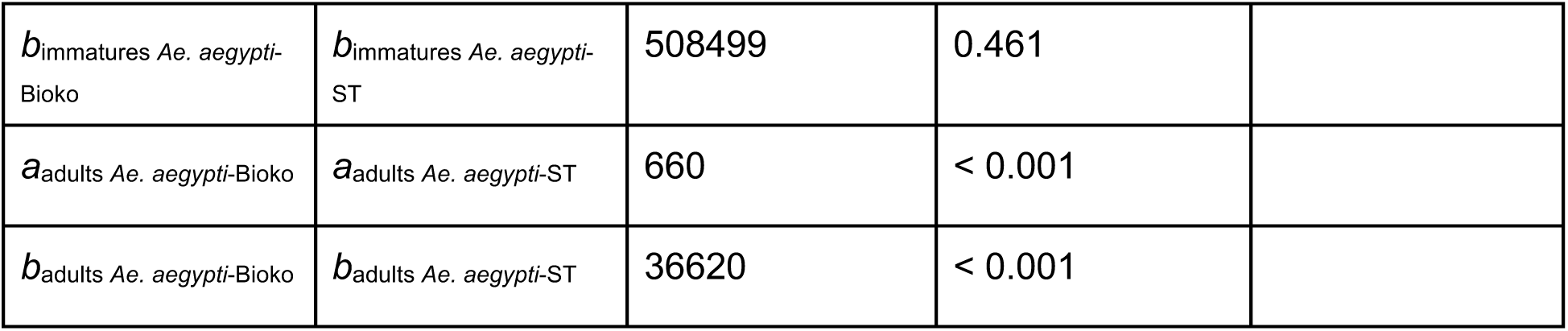
Pairwise comparisons between parameters of the exponential functions. *a*: Intercept, b: rate of decay.

| Parameter 1 | Parameter 2 | W | P-value |  |
| --- | --- | --- | --- | --- |
| $a_{\text{immatures Ae. albopictus-Bioko}}$ | $a_{\text{adults Ae. albopictus-Bioko}}$ | 629898 | < 0.001 | |
| $b_{\text{immatures Ae. albopictus}}$ | $b_{\text{adults Ae. albopictus}}$ | 995859 | < 0.001 | |
| $a_{\text{immatures Ae. aegypti-Bioko}}$ | $a_{\text{adults Ae. aegypti-Bioko}}$ | 990211 | < 0.001 | |
| $b_{\text{immatures Ae. aegypti-Bioko}}$ | $b_{\text{adults Ae. aegypti-Bioko}}$ | 996602 | < 0.001 | |
| $a_{\text{immatures Ae. albopictus-ST}}$ | $a_{\text{adults Ae. albopictus-ST}}$ | 36331 | < 0.001 | |
| $b_{\text{immatures Ae. albopictus-ST}}$ | $b_{\text{adults Ae. albopictus-ST}}$ | 163405 | < 0.001 | |
| $a_{\text{immatures Ae. albopictus-Bioko}}$ | $a_{\text{immatures Ae. albopictus-ST}}$ | 62869 | < 0.001 | |
| $b_{\text{immatures Ae. albopictus-Bioko}}$ | $b_{\text{immatures Ae. albopictus-ST}}$ | 4520 | < 0.001 | |
| $a_{\text{immatures Ae. aegypti-Bioko}}$ | $a_{\text{immatures Ae. aegypti-ST}}$ | 499329 | < 0.001 | |
| *b*<sub>immatures Ae. aegypti-Bioko</sub> | *b*<sub>immatures Ae. aegypti-ST</sub> | 508499 | 0.461 |  |
| *a*<sub>adults Ae. aegypti-Bioko</sub> | *a*<sub>adults Ae. aegypti-ST</sub> | 660 | < 0.001 |  |
| *b*<sub>adults Ae. aegypti-Bioko</sub> | *b*<sub>adults Ae. aegypti-ST</sub> | 36620 | < 0.001 |  |

